# Differential and overlapping effects between exogenous and endogenous attention shape perceptual facilitation during visual processing

**DOI:** 10.1101/2022.12.23.521777

**Authors:** Mathieu Landry, Jason Da Silva Castanheira, Karim Jerbi

## Abstract

Visuospatial attention is not a monolithic process and can be divided into different functional systems. In this framework, exogenous attention reflects the involuntary orienting of attention resources following a salient event, whereas endogenous attention corresponds to voluntary orienting based on the goals and intentions of individuals. Previous work shows that these attention processes map onto distinct functional systems, yet evidence suggests that they are not fully independent. In the current work, we investigated the differential and overlapping effects of exogenous and endogenous attention on visual processing. We combined spatial cueing of visuospatial attention, electroencephalography (EEG), and multivariate pattern analysis (MVPA) to examine where and when the effects of exogenous and endogenous attention were maximally different and maximally similar. Critically, MVPA provided new insights by examining whether classifiers trained to decode the cueing effect for one attention process (e.g., exogenous attention) can successfully decode the cueing effect for the other attention process (e.g., endogenous attention). These analyses uncovered differential and overlapping effects between exogenous and endogenous attention. Next, we combined principal component analyses, single-trial event-related potentials, and mediation analysis to determine whether these effects facilitate perception, as indexed by the behavioral spatial cueing effects of exogenous and endogenous attention. This approach revealed that three EEG components shape the cueing effects of exogenous and endogenous attention at various times after target onset. Altogether, our study provides a comprehensive account about how overlapping and differential processes of endogenous and exogenous relate to perceptual facilitation in the context of visuospatial attention.

**Significance Statement:** Top-down and bottom-up attention represent separate functional systems in the brain. Previous research suggests, however, that they are not fully independent, and can interfere with each other. In the present study, the authors use machine learning techniques and recordings of brain activity to investigate differences and similarities between top-down and bottom-up attention during the visual processing of stimuli. This approach allowed them to explore how top-down and bottom-up attention processes facilitate perception. Their results show that top-down and bottom-up attention operate differently as early as 100ms after the onset of a target. In contrast, they operate similarly 200ms after the target onset. Most importantly, these effects are directly related to the participants’ perceptual behavior. In sum, our study shows that top-down and bottom-up attention support the perception of stimuli through overlapping and distinct spatio-temporal brain patterns.

## Introduction

Prevailing views differentiate top-down and bottom-up attention processes (Knudsen, 2007). In this framework, exogenous attention (i.e., bottom-up attention) reflects involuntary and stimulus-driven orienting of attention resources based on the features of a stimulus (e.g., your phone’s alarm going off). On the other hand, endogenous attention (i.e., top-down attention) corresponds to the voluntary control of attention resources based on the goals and intentions of individuals (e.g., reading a book (Carrasco, 2011). Several lines of research emphasize that exogenous and endogenous map onto distinct brain processes, which supports the theory that both attention systems operate through independent and distinct processes (Chica, Bartolomeo, & Lupiáñez, 2013).

A significant part of research on these attention processes follows from the spatial cueing paradigm – an experimental approach designed to orient attention to a spatial location using a cue (Chica, Martín-Arévalo, Botta, & Lupiánez, 2014). A target stimulus occurs at different spatial locations, while participants are asked to report on (i.e., detect, locate, discriminate, or identify) said target stimulus. Here, the effects of attention are typically evaluated by comparing trials where the target event occurs at a cued location versus trials where the target event occurs at an uncued location. Hence, the comparison of cued and uncued trials reveal the perceptual benefits of visuospatial attention – i.e., the spatial cueing effect – wherein participants are quicker and more accurate to respond for target stimuli occurring at the attended locations versus unattended locations. In this experimental approach, exogenous attention is often engaged via a task-irrelevant peripheral salient cue that prompts a stimulus-driven response (Posner, 1980), whereas endogenous is engaged via a task-relevant central cue indicating to participants where to voluntary orient their attention (Jonides, 1981).

Research in neurophysiology shows that visuospatial attention hardly reduces to a singular neural process, but instead comprises several stages of processing (Malkinson et al., 2022; Martín-Arévalo, Chica, & Lupiáñez, 2016). In this regard, an important body of work highlights the benefits of exogenous and endogenous attention 100ms after the target onset. Both attention systems alter the P1 and N1 event-related potentials (ERP) at the contralateral occipital region (for review, see Luck, Woodman, & Vogel, 2000; also, Mangun, 1995). These early effects of attention along the visual stream are thoughts to reflect sensory gains, which results in perceptual facilitation (Dosher & Lu, 2000; Hillyard, Vogel, & Luck, 1998; Itthipuripat, Ester, Deering, & Serences, 2014). However, while both exogenous and endogenous attention modulate early sensory processes, mounting evidence suggests that they may do so differently. Using a double cueing approach where both exogenous and endogenous attention are concurrently engaged, Hopfinger and West (2006) uncovered an interaction between both attention systems over the N1 component. This interaction implies differential visual processing effects of exogenous and endogenous attention. This outcome is consistent with results from brain imaging (e.g., Dugué, Merriam, Heeger, & Carrasco, 2020). Our recent EEG work aligns with these findings and showed that exogenous and endogenous attention interfere with each other early during visual processing (Landry, da Silva Castanheira, Raz, Baillet, & Sackur, 2022).

Endogenous and exogenous attention also both modulate later stages of visual processing, like the P2 (e.g., McDonald, Ward, & Kiehl, 1999), N2 (e.g., Mangun, Hillyard, & Luck, 1993) and P3 (e.g., Chica & Lupiáñez, 2009) ERPs components. These effects are often thought to indicate target-related processing such as feature selection, the filtering of task-irrelevant events, and the accumulation of perceptual evidence for target discrimination (Akyürek & Schubö, 2013; Luck & Hillyard, 1994; Nunez, Vandekerckhove, & Srinivasan, 2017). Neurophysiological findings similarly suggest differential effects between exogenous and endogenous attention for these later stages of target processing. Previous work showed that endogenous attention increases the amplitude of the N2 and P3 components, while researchers observed no such effect for exogenous attention (Hopfinger & West, 2006; Wang, Wu, Fu, & Luo, 2010).

Although previous EEG work highlights distinct and shared processes between endogenous and exogenous attention along the visual stream, it remains uncertain how these effects contribute to behavior. The goals of the present study were threefold. First, we aimed to corroborate previous findings suggesting the differential effects of exogenous and endogenous on target processing. We trained multivariate pattern analysis (MVPA) to classify the effects of exogenous and endogenous attention and extracted when and where the attention systems were maximally different. Second, we aimed to investigate overlapping neural processes between exogenous and endogenous attention. For this second goal, we examined whether classifiers trained to decode the cueing effect for one form of attention processing generalizes to the other form of attention processing (e.g., test the performance of classifiers trained to decode exogenous cue validity in the context of endogenous cue validity). Based on previous work, we hypothesized that exogenous and endogenous attention would differ early during the processing of sensory events, around the P1-N1 complex, and would be similar further down the visual stream. Our third and last objective was to relate neurophysiological processes of endogenous and exogenous attention to behaviour. We aimed to determine whether distinct and shared processes of attention contribute to the exogenous and endogenous cueing effects. We relied on single-trial ERPs to breakdown visuospatial attention into neurophysiological components and used a mediation analysis to test whether these components contribute to the facilitation attention effects observed at the behavioral level. Altogether, our findings highlight where and when the brain processes of exogenous and endogenous attention were different and similar, and whether these differences and similarities contribute to the observed behavioral facilitation effects of attention.

## Methods

### Participants

We recruited 42 participants based on convenience sampling. Participants received monetary compensation of $10/hour for completing 4 different tasks. Each task consisted of 384 trials. Participants performed 10 practice trials before each task. We analyzed data from 3 tasks: the non-cueing task, the endogenous cueing task, and the exogenous cueing task. Data from all four tasks has been disseminated elsewhere (Landry et al., 2022). The experiment was a single testing session and lasted approximately 2 hours. All participants had normal or corrected-to-normal vision and provided consent. The experimental protocol was approved by the McGill Ethics Board Office.

Seven participants were excluded due to poor EEG data quality. One participant did not complete all tasks. We additionally excluded one participant due to below chance-level discrimination accuracy rate (∼40% averaged across all conditions) and one participant due to a high volume of timeout errors (response times > 1500ms on ∼11% of trials overall). The final sample included 32 individuals (24 women, Mean age = 21.9 (SD = 2.6)). We used G*Power3 (Faul, Erdfelder, Lang, & Buchner, 2007) to perform sample size calculations across exogenous and endogenous cueing tasks based on effect size estimates reported in the review of Chica et al. (2014). For repeated measures F-test in the context of a central predictive cue and long cue-target latencies (i.e., > 500ms), we calculated that 6 participants are required to achieve a power of .8 based on a large effect size (η^2^ = .34) and alpha of .5. Likewise, for a peripheral non-predictive cue and short cue-target latencies (i.e., >300ms), 6 participants are also required to achieve a power of .8 based on a large effect size (η^2^ = .39) and alpha of .5. The sample size of the current study is therefore more than adequate to detect the effects of exogenous and endogenous attention at the behavioral level. Furthermore, note that our sample size is almost twice that of previous EEG experiments investigating exogenous and endogenous orienting together (Hopfinger & West, 2006; Keefe & Störmer, 2021).

### Stimuli, Apparatus & Design

Participants viewed stimuli on a 24-in BenQ G2420HD monitor sitting approximately 60 cm away. Stimulus presentation was done using MATLAB R2015b (Mathworks Inc., Natick, MA, USA) and the third version of the Psychophysics toolbox (Brainard, 1997; Kleiner et al., 2007; Pelli, 1997). The screen was set to 75hz. Except for the target, all stimuli were black (i.e., RGB values of 0, 0, 0; 1.11 cd/m^2^) and white (i.e., RGB values of 255, 255, 255; 218.8 cd/m^2^) drawings on a grey background (i.e., RGB values of 128, 128, 128; 70.88 cd/m^2^). The fixation marker was an empty circle made from a black line drawing with a radius of 1.2° located in the center of the screen. Two target placeholders were located at 8.7° on each side of the fixation marker on the left and right side of the screen. These placeholders were made from black line drawings of empty circles with a 2.4° radius. We cued exogenous attention by briefly changing the line drawing from one of the placeholders to white. To ensure that this cue solely engaged exogenous orienting, the cue-target spatial contingency was set to 50%, such that the cue was only predictive of the target’s location at chance-level. The exogenous cue was therefore non-informative and occurred at the periphery. This is consistent with the standard approach for eliciting exogenous attention in the lab (Chica et al., 2014). We cued endogenous attention by coloring the inside of the fixation marker, wherein the right or left half of the circle was shaded in black, and the other half in white. The side of the fixation marker that turned white indicated where the target was likely to occur. For example, if the right side turned white, the target was 66.6% likely to appear in the right placeholder. In this way, we avoided using overlearned directional cues, like an arrow, to engage endogenous attention (Ristic & Landry, 2015). Participants were aware of these contingencies. The targets were sinusoidal black and white gratings combined with a Gaussian envelope. Spatial frequency was set to 3 cpd. Target stimuli were tilted 5° degrees clockwise or counterclockwise.

### Procedure

Participants completed four tasks of 384 trials: A non-cueing task, an exogenous cueing task, an endogenous cueing task, and a double cueing task where both exogenous and endogenous orienting were engaged. Task order was randomized across participants. Note that the current manuscript only includes data from the non-cueing, the endogenous cueing, and the double cueing tasks.

Participants were encouraged to maintain their fixation at the center of the screen throughout the experiment, while we assessed their eye movements using electro-oculogram. We jittered the latencies between fixation and attention cues, between both attention cues, and between the cues and the target. We used a uniform distribution of latencies to minimize the effects of temporal prediction following spatial cueing. Cue-target latencies were adjusted following the temporal profiles of exogenous and endogenous attention so that the benefit of attention processing would be optimized for both attention (Chica et al., 2014).

In the non-cueing task, the timing between the fixation circle and target stimulus was jittered from 1027 to 1280ms. The target stimulus stayed on the screen until participants responded. We used the non-cueing task as a baseline against which we regressed the sensory effects of cue stimuli for the target-related analysis (see the *Electroencephalography* section below). We also used the target-locked ERP from the non-cueing task to extract EEG components pertaining to visual processing in the absence of spatial cueing of exogenous and endogenous attention.

In the exogenous orienting task, again the timing between the fixation circle and target stimulus was jittered between 1027 and 1280ms, followed by the onset and offset of the exogenous cue. The onset of the exogenous cue was jittered from 106 to 307ms before target onset. The exogenous cue always stayed 106ms on screen before offsetting. The timing between the onset of the endogenous cue and target onset was jittered between 613ms and 926ms. The target stimulus remained on screen until participants responded (Figure 1). In the endogenous orienting task, again the timing between the fixation circle and target stimulus was jittered between 1027ms and 1280ms, followed by the endogenous cue that remained on the screen until participants responded. The timing between the onset of the endogenous cue and target onset was jittered between 613ms and 926ms. The target stimulus remained on screen until participants responded (Figure 1). Participants were instructed to complete a target discrimination task and indicate the orientation of the Gabor target as quickly and accurately as possible on a QWERTY keyboard by pressing the F key for counterclockwise orientation and the J key for clockwise orientation. Inter-trial period was set to 1s.

**Figure 1.**
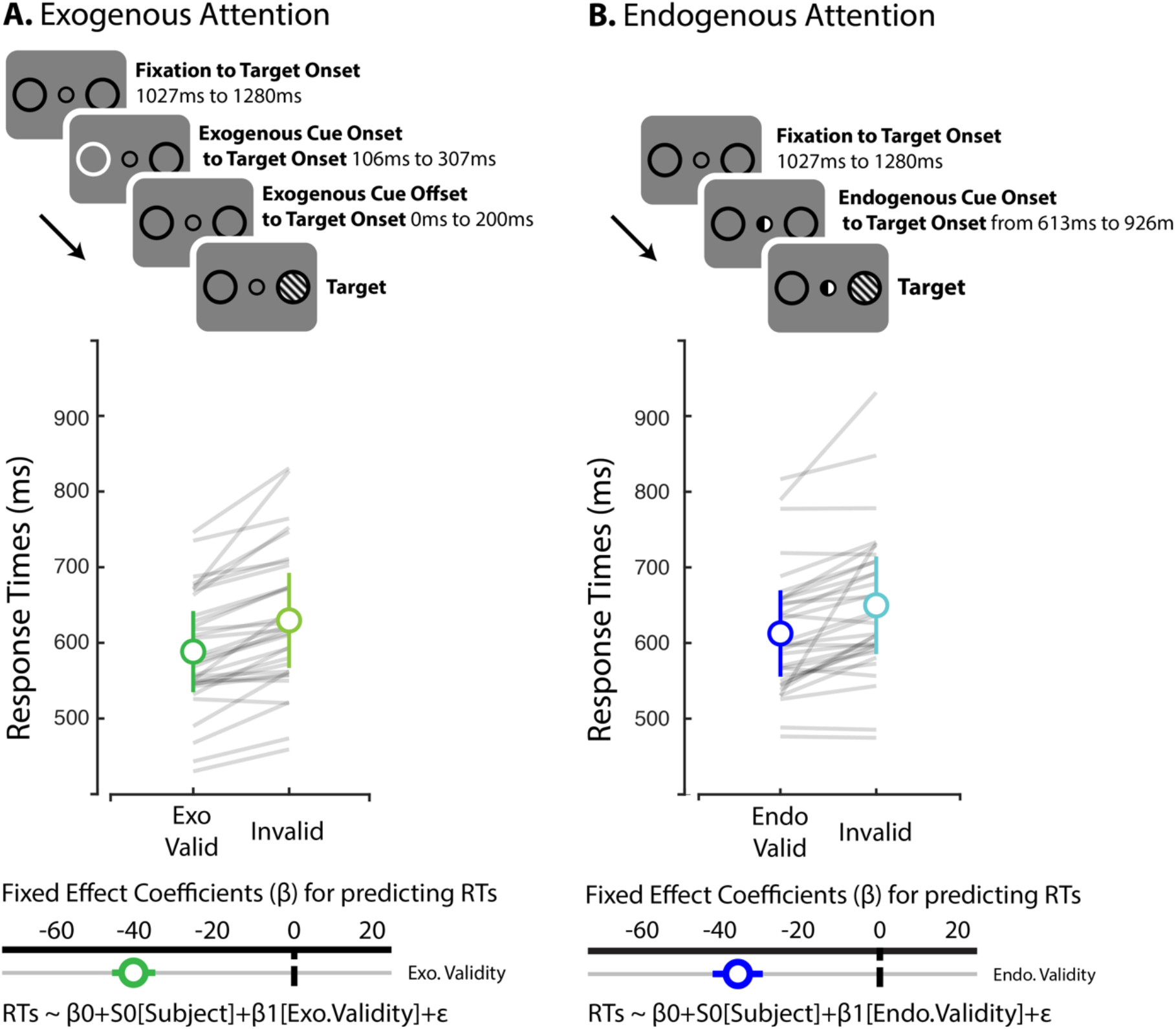
**A.** For the exogenous spatial cueing task, each trial began with a fixation screen, followed by the onset of the exogenous cue (i.e., a target placeholder would briefly flash) after a random interval. The cue would be offset after 200ms. Lastly, a Gabor stimulus was shown at the left or right target location until the participant’s discrimination response. The exogenous cue was non-predictive of the target’s location. **B.** For the endogenous spatial cueing task, each trial began with a fixation screen followed by the onset of the endogenous cue (i.e., half of the black fixation circle would become white) after a random interval. The cue stayed on the screen until the end of the trial. The endogenous cue was task-relevant and predicted the target’s location. In both cueing tasks the onsets of the endogenous cue, the exogenous cue, and the target stimulus were time jittered. The plots show average (black dots) and participants (color dots) accurate RTs across conditions. Error bars represent bootstrapped 95% C.I. Bottom graphs indicate hierarchical regression model coefficients and corresponding 95% C.I for evaluating cue validity.

### Data analyses for behavioral performance

Participants’ discrimination performance was near ceiling; the average accuracy rate was ∼93%. Thus, we examined task performance via accurate response times (RTs). We removed trials where participants made anticipation (i.e., RTs < 150ms) or timeout (i.e., RTs > 1500ms) errors. This accounted for less than 2% of trials overall. We additionally removed wrong key presses, which corresponded to less than 1% of trials. We used hierarchical linear regression models (Gelman & Hill, 2006) to test the effects of exogenous and endogenous attention on task performance, where we included cue validity as a fixed factor and participants as a random one:

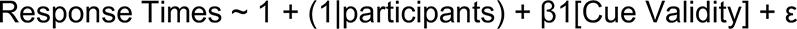

To compare the cueing effects between exogenous and endogenous attention, we included cue validity (i.e., valid versus invalid) and task (i.e., exogenous cueing and endogenous cueing) as fixed factor, and participants as a random one.

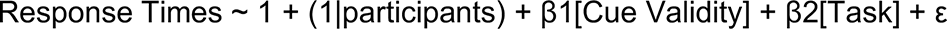

Fixed factors were added in stepwise fashion while we used a chi-square goodness-of-fit test over the deviance to determine whether they significantly improved the fit. We computed the Bayesian information criterion (BIC) to select the most parsimonious model. Lastly, we calculated Bayes factors to weight evidence for the alternative against the null hypothesis based on the BIC approximation (Wagenmakers, 2007) :

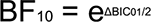

### Electroencephalography

We recorded EEG signals using 64 Ag/AgCl active electrodes at a sampling rate of 1000 Hz (ActiCap System; Brain Products GmbH; Gilching, Germany). We monitored eye blinks and eye movements through additional bipolar electrodes placed at the outer canthi, as well as the superior and inferior orbits of the left eye. We kept impedances of all electrodes below 10 kΩ, while all electrophysiological signals were amplified (ActiChamp System; Brain Products GmbH; Gilching, Germany). Electrodes were referenced online to C4. We re-referenced the electrodes offline to the average of all channels. Preprocessing and analysis were conducted in BrainVision Analyzer (ActiChamp System; Brain Products GmbH Inc.; Gilching, Germany) and MATLAB (R2020a; Mathworks Inc., Natick, MA) using Brainstorm (Tadel, Baillet, Mosher, Pantazis, & Leahy, 2011) and custom scripts. We downsampled the data to 250 Hz and visually inspected EEG signals to identify activity exceeding ± 200 μV. We applied two IIR Butterworth filters: a first High-pass 0.1 Hz filter of 8th order and a 60Hz notch filter. We interpolated bad channels topographically (1.12% of channels) and then identified artifacts related to eye movements and blinks using independent component analysis via the BrainVision Analyzer Ocular correction ICA tool. Despite our conventional and robust approach to remove eye movement artefacts from the EEG, residuals from this procedure could have influenced our MVPA analyses for examining differential and overlapping effects between exogenous and endogenous attention. However, evidence strongly argues against this possibility. First, we examined eye movements based on HEOG and VEOG channels as a function of exogenous and endogenous attention for leftward and rightward orienting of attention. We found no significant difference between exogenous and endogenous based on pairwise t-tests where we compared the time course of these ocular channels. Second, the classifiers’ weights from all MPVA emphasized the importance of the posterior region of the scalp for all classification problems. If classification performance was driven by eye movements artifacts, the MPVA weights would instead indicate anterior parts of the scalp. Third, our findings from the MVPA were also consistent with the outcomes of our single-trial ERP mediation analysis, whereby we observed strong effects of exogenous and endogenous attention on components related to visual processing that were extracted from a non-cue condition – these visual components are therefore distinct from eye movements. Lastly, the results from our MVPA also aligned with previous findings in the field regarding early sensory processing.

The close temporal proximity of the cue and target stimuli entails that activity from the former may overlap with that of the latter. To remove this overlapping activity, instead of using the adjacency response technique (Woldorff, 1993), we removed event-related responses from exogenous and endogenous cue stimuli using regression modelling. Here we took the residuals of the linear regression model using the non-cueing condition as baseline. A hierarchical linear regression was run in MATLAB with cueing condition as a dummy coded fixed factor, and subjects as a random factor. ‘Raw’ residuals were obtained from all channels and subjects.

### Event-related potentials

We analyzed target-related event-related potentials (ERPs). We first applied a FIR bandpass-pass filter between 0.5 and 15 Hz and then divided the EEG into epochs spanning −200 and 1000ms. All triggers were realigned according to a photodiode stimulation linked to the onset of the target event. ERPs were baseline corrected from -100 to 0ms.

### Multivariate analyses

We leveraged multivariate statistical techniques to evaluate the influence of exogenous attention on the cueing effects of endogenous attention based on target-locked ERP. Our approach was twofold. Our first goal was to uncover the differential effects between exogenous and endogenous attention using MVPA (Figure 2A). Here, we tested whether SVM classifiers can differentiate exogenous cue validity (exogenous cue valid ERP minus exogenous cue invalid ERP) from endogenous cue validity (endogenous cue valid ERP minus endogenous cue invalid ERP). Our second goal aimed to uncover overlapping effects between them by training classifier to separate cue validity for one attention process (e.g., exogenous cue valid ERP versus exogenous cue invalid ERP) and then testing them in the context of the other attention process (e.g., endogenous cue valid ERP versus endogenous cue invalid ERP), and vice-versa (Figure 2B).

**Figure 2.**
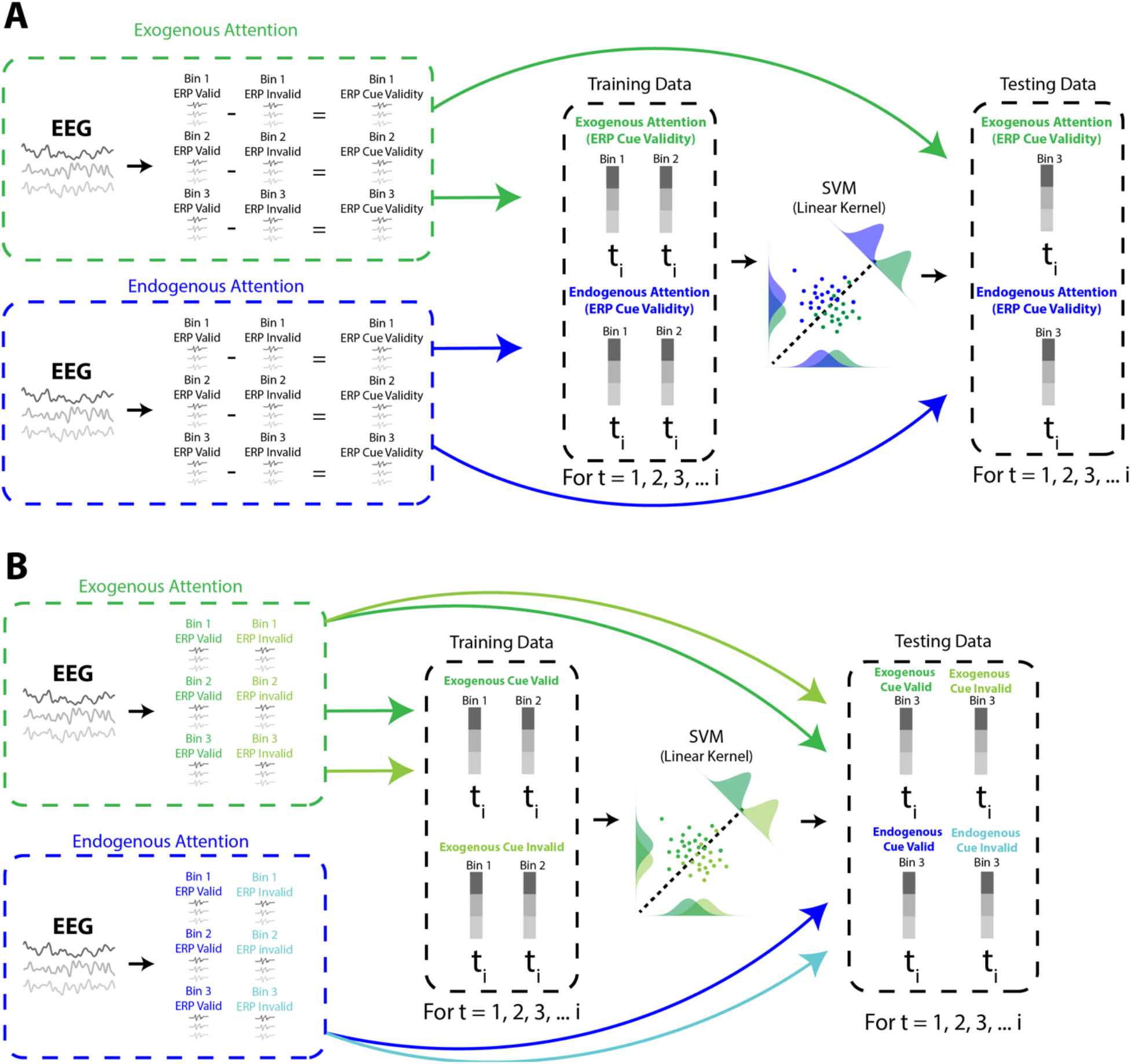
Decoding pipeline **A**. For each participant, across all channels, we randomly separated epoched EEG time series into three bins for cue valid trials and cue invalid trials separately, then averaged EEG to produce an ERP per bin, and lastly subtracted cue invalid ERP from cue valid ERP for each corresponding bins. We perform these procedures in parallel for exogenous and endogenous attention. We used SVM to decode exogenous from endogenous attention. For each time point, we trained the classifier using two data points for each form of attention, and then tested it on the remaining one. We iterated this process three times so that each averaged epoch was used twice for training and once for testing. We repeated this process 50 times per participant. **B**. Using a similar approach, we trained classifiers to decode cue validity for each attention process separately (e.g., exogenous cue valid ERP vs. exogenous cue invalid ERP) and then tested whether the trained SVM model for each time point generalizes to the other attention process (e.g., whether SVM models trained to decode exogenous cue validity generalize to endogenous cue validity).

Our multivariate statistical analyses largely follow from the work of Bae and Luck (2018). Here, we performed multivariate classification using linear support vector machine (SVM) and MATLAB’s *fitcsvm* and *predict* functions. The training, validation and testing phases were completed at the participant’s level. As we mentioned previously, our first analysis aimed to differentiate exogenous cue validity ERP from endogenous cue validity ERP where we used a three-fold cross-validation procedure to train the classifier on 2/3 of the trials and then validated it on the remaining 1/3. Following this three-fold cross-validation approach, target-locked EEG from trials were separated into three bins per cue validity for each attention cueing tasks separately – i.e., three bins of trials for exogenous cue valid and three bins for exogenous cue invalid, as well as three bins of trials for endogenous cue valid trials and three bin for endogenous cue invalid trials. We equated trials across all bins for all participants and all conditions. There were 28 trials per bin. We extracted EEG derivations from each channel as a function of ipsi- and contralateral location relative to target location for each trial. We then averaged the trials from each bin and baseline corrected them from −100ms to 0ms, which resulted in three separate target-locked waveforms for cue valid trials and three separate waveforms for cue invalid trials for all 64 EEG channels across both exogenous and endogenous attention cueing tasks. Next, for each attention cueing condition and each channel separately, we subtracted the waveform of the first bin for cue invalid trials from the waveform of the first bin for cue invalid trials, and then the same for the second bin and lastly for the third bin. This procedure resulted in three waveforms that corresponded to the exogenous cue validity effect, and three waveforms that corresponded to the endogenous cue validity effect. Then, we applied SVM for each time point along the time series by including the 64 channels as features in our model. We trained the classifier using two waveforms from each class (i.e., two waveforms for exogenous attention and two for endogenous attention and then validated it using the remaining ones (i.e., one waveform for exogenous attention and one for endogenous attention). We performed permutations between the training and validation phases, whereby each EEG time series was used twice for training and once for testing. We repeated this process 50 times per participant while randomly shuffling trials across bins for each repetition. Lastly, we averaged classification accuracy rates for each participant across both target locations and smoothed these values across the time series via a five-sample sliding window -- i.e., a 20ms window (Hong, Bo, Meyyappan, Tong, & Ding, 2020). This procedure allowed us to assess the classification performance across the time series during single cueing. Figure 2A provides a diagram of the overall procedure.

We adopted the same approach to achieve our second objective and explore overlapping effects between exogenous and endogenous attention. However, in this instance, we wanted to train classifiers to separate cue valid trials and cue invalid trials for one attention process and assess whether classification generalizes to the other attention process. In this instance, for exogenous attention, we separated target-locked EEG derivations from each trial into three bins of trials for cue invalid trials and three bins for valid trials. Again, there were 28 trials per bin. This was done for each EEG channel separately. We then averaged the time-series in each bin, which resulted in three target-locked waveforms for exogenous cue valid trials and three waveforms for exogenous cue invalid trials. In this process, we also separately binned 28 trials from the endogenous cue valid condition and 28 from the endogenous cue invalid condition and averaged the EEG time series to obtain target-locked waveforms that were baseline corrected from - 100 to 0ms. SVM classifiers were trained and validated to correctly classify exogenous cue valid waveforms and exogenous cue invalid waveforms across the time series. Additionally, we tested the models that were trained on exogenous cue valid to correctly classify endogenous cue valid waveform and endogenous cue invalid waveform. Lastly, we applied the same procedure to train and validate our model on endogenous attention, and then testing it on exogenous attention. Figure 2B provides a diagram of the overall procedure

We used one sample t-tests across the time series to determine whether classification performances were better than the analytical chance-level (i.e., 50%). We also compared classification performance between decoding of exogenous and endogenous attention using pairwise t-tests. We controlled for family-wise errors via cluster-corrected mass permutation t-tests. The cluster forming threshold was set to p < .05. We performed 1000 permutations where we randomly varied the classification labels and then contrasted observed cluster sizes based on t-statistics against surrogate distributions. The threshold for statistically significant clusters was set to 95%.

### Single-trial ERP

We aimed to determine whether differential and overlapping effects between exogenous and endogenous attention contribute to perceptual facilitation. To investigate this question, we combined single-trials ERP analysis with simple mediation modelling. We aimed to determine whether modulations of ERP amplitude mediate the relationship between spatial cueing of attention and accurate response times. Here, we performed separate analyses for exogenous and endogenous attention.

To perform single-trial ERP, we first used baseline corrected (−100 to 0ms to target onset) target-locked ERP from the non-cueing task to isolate components related to visual processing independently from exogenous and endogenous attention processes. We calculated averaged waveforms for each channel as a function of target location – i.e., ipsi- and contralateral processing. We then applied PCA with EEG channels as features and the time series as observations to these waveforms (i.e., applying PCA to time series by channels matrix). Our goal was to uncover the linear combination of channels that explain the most variance for non-cueing (i.e., neutral) target-locked ERP. To uncover the most important components, we performed 1000 permutations where we randomly varied channel location and extracted the total variance explained for each iteration. We then established the 95% threshold based on this null distribution of explained variance (Cohen, 2014). This approach isolated three components. The loading coefficients of the first component revealed a posterior ipsilateral topographical pattern (see Appendix B). Projecting the averaged target-locked waveforms onto this first component revealed P1 and N1 at 132ms and 240ms post-target onset, respectively, as well as a P2 at 356ms (see Appendix C). The loading coefficients of the second component showed a posterior contralateral topographical pattern (see Appendix B). Projecting the averaged target-locked waveforms onto this second component revealed P1 and N1 at 104ms and 164ms post-target onset, as well as a P3 that peaked at 340ms (see Appendix C). The loading coefficients of the third component showed a posterior contralateral topographical pattern (see Appendix B). Projecting the averaged target-locked waveforms onto this third component revealed P1 and N1 at 84ms and 136ms post-target onset, and the N2 that peaked at 304ms (see Appendix C).

Next, we projected each trial from the exogenous and endogenous cueing tasks onto all three latent components to get single-trial ERP, whereby the loading coefficients (i.e., vector of loading coefficients) weighted the data of each trial (i.e., time series by channels; Appendix C). This approach improved the signal-to-noise ratio while we explore the effects of exogenous and endogenous attention across three latent variables (Nunez et al., 2017). In this way, we obtained single-trial ERP across all three components for each attention process. Appendix C shows the averaged waveforms for all trials and participants across all 3 components for exogenous and endogenous attention cue validity.

### Mediation Analysis

Using single-trial ERP, we then tested our hypothesis that early differential and late overlapping effects between exogenous and endogenous attention contribute to perceptual facilitation using meditation analyses. We relied on hierarchical regression models and the Sobel approach to estimate the indirect effect via the product of regression coefficients (Sobel, 1982). We used on the following regression models to estimate our indirect effect, wherein cue validity and the amplitude of the ERP were used as fixed factors, while participants were added as a random factor:

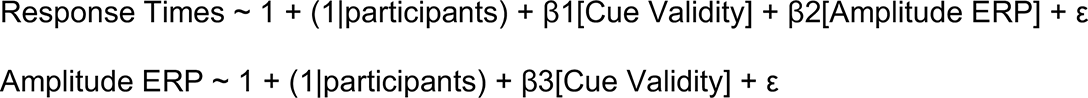

We looked at the mediation coefficients across the entire time series. Here, we calculated a z-score and p-value for each time point. This approach allowed us to use cluster-corrected mass permutation t-tests. Again, the cluster forming threshold was set to p < .05. We performed 1000 permutations where we randomly varied the cue validity variable and RTs. Lastly, we contrasted observed cluster sizes based on t-statistics against surrogate distributions. The threshold for statistically significant clusters was set to 95%. We applied this approach across all three components and attention processes separately. We calculated the variance explained related to each significant cluster of the indirect effect using the following equation (Fairchild, MacKinnon, Taborga, & Taylor, 2009):

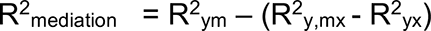

## Results

We collected data from thirty-two participants who completed three tasks while recorded EEG: An endogenous cueing task, an exogenous cueing task, and a no cueing task. Across all tasks, participants were asked to discriminate the orientation of a Gabor target (clockwise versus counterclockwise). The target event occurred at a cued or uncued location during spatial cueing (Figure 1).

### Behavior

Discrimination performance was near the ceiling (∼93% average accuracy rate overall). We therefore focused our behavioral analysis on accurate response times (RTs). Averaged RTs for the non-cued condition was 610.02ms 95% CI [581.64, 644.68]. Next, we confirmed that the exogenous and endogenous cues yielded facilitation effects. We used hierarchical regression models with cue validity (cue valid vs. cue invalid) as a fixed factor and participants as a random one. This analysis shows that participants were ∼41ms faster for accurately discriminating the orientation of Gabor target for cued trials for exogenous cueing (exogenous cue validity: β=-41.28, SE=2.82, 95% CI [−46.83, −35.74]; Figure 1A). Participants were likewise ∼36ms faster to discriminate Gabor targets for cued trials for endogenous attention (endogenous cue validity effect; β=-36.49, SE=3.24, 95% CI [−42.84, −30.13]; Figure 1B).

Having confirmed perceptual facilitation across both spatial cueing tasks, we tested whether exogenous and endogenous cue validity effects differed. Again, we used hierarchical regression modelling where we included cue validity (cue valid vs. cue invalid), task (exogenous cueing task vs. endogenous cueing task), and their interaction as fixed factors and participant as a random factor. The interaction between cue validity and task examined whether cue validity differed between exogenous and endogenous cueing tasks. We added predictors in a stepwise fashion. The best fitting regression model revealed main effects of cue validity (β=-38.87, SE=2.17, 95% CI [−43.13, −34.61]) and task (β=21.13, SE=2.14, 95% CI [16.93, 25.33]). Hence, participants were on average ∼38ms faster for cued versus uncued trials across both tasks, while the main effect of task revealed that they were overall ∼21ms slower during the endogenous cueing task compared to exogenous task. Critically, when we tested the interaction between cue validity and task, we confirmed that the cue validity effect did not differ between exogenous and endogenous attention (β=5.07, SE=4.36, 95% CI [−3.46, −13.61]). Evidence strongly weighted against the likelihood of the interaction between cue validity and task variables (BF01 = 76.55).

### Electrophysiology: Decoding differential effects of exogenous and endogenous attention

Using a MVPA approach adapted from Bae and Luck (2018), we tested where and when exogenous and endogenous cue validity effects differed along the target-locked EEG time series. We trained linear support vector machine (SVM) classifiers for each timepoint and subject, independently. We randomly split trials into three equally sized bins and averaged the time series of EEG potentials within each bin, yielding an ERP for each bin (see Figure 2A). We did this separately for cue valid trials and cue invalid trials, and then subtracted the ERP of cue valid trials from the ERP of cue invalid trials across corresponding bins. We applied this procedure separately for exogenous and endogenous attention. This process produced 6 separate time series: 3 distinct time-series reflecting the exogenous attention cue validity effect and 3 time-series reflecting the endogenous attention cue validity effect. We used a three-fold cross-validation strategy to train classifiers to decode which attention process is engaged (i.e., endogenous vs exogenous attention). Here, two of the three cue validity time-series for each attention system (i.e., four in total) were used to train classifiers, and the remaining two (i.e., one per attention system) were used to test the classifiers’ performance. We iterated across all possible combinations such that each time series was used twice for training and once for testing. The respective decoding accuracies of the classifiers were determined from their correct classification performance on the left-out time-series (i.e., the test data set). We repeated the above random binning procedure 50 times per participant (see Methods and Figure 2A). We averaged decoding accuracies across all iterations and used pairwise mass t-statistics across the time series to evaluate accuracy rates against chance-level.

This analysis supported our primary hypothesis about differential effects. We observed that classification performance for differentiating exogenous and endogenous cue validity was better than chance-level early after target onset – i.e., 20ms post-target stimulus (Figure 3A). Decoding accuracy for separating exogenous and endogenous cue validity effects was above chance-level between 20ms until 800ms. The average maximal decoding performance across participants occurred at 172ms (95% CI, [143ms 201ms]) after target onset (Figure 3B). We observed a contralateral posterior fluctuation of SVM weights (i.e., the coefficients of the classifiers) at 172ms (Figure 3C). Altogether, these results highlight that exogenous and endogenous attention produce different cue validity effects early during sensory processing (i.e., starting at ∼20ms).

**Figure 3.**
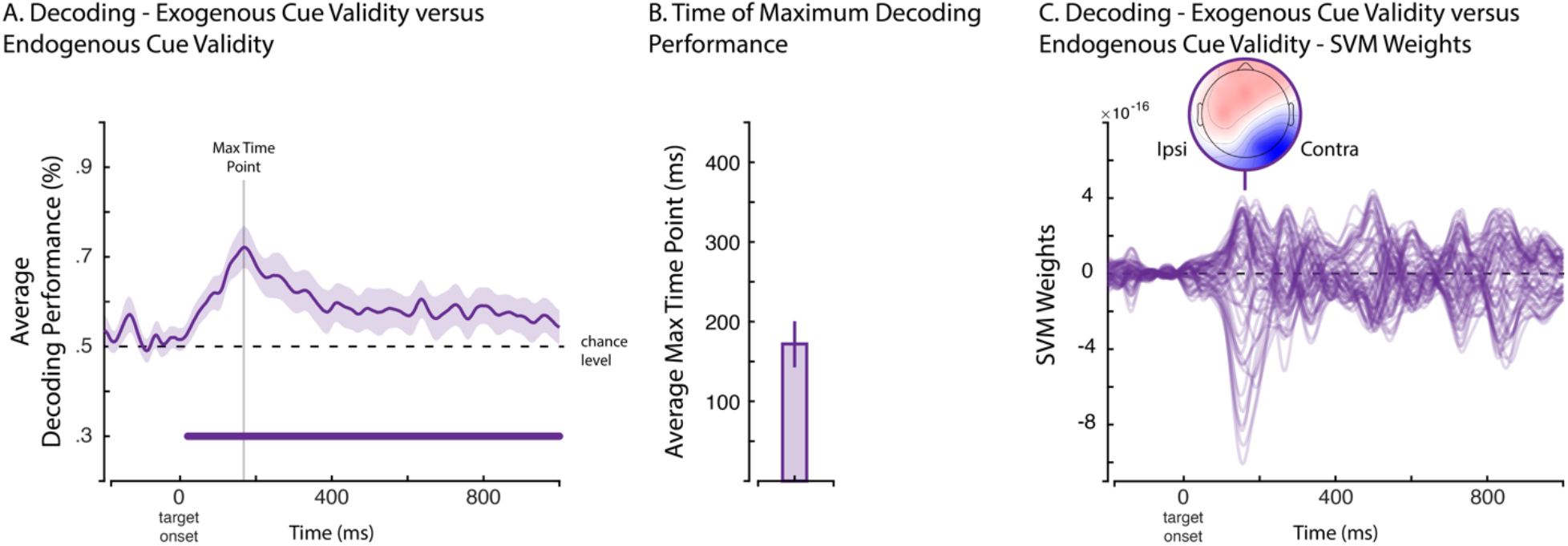
**A**. Group averages of decoding accuracy from target-locked EEG traces for exogenous cue validity versus endogenous cue validity. We used one sample t-tests and random cluster permutation to test for significant differences in classification performances against chance-level (50%). Horizontal purple line indicates the temporal segments of significant differences in decoding accuracy. Shaded areas represent the 95% C.I. **B**. Averaged maximal time point for decoding accuracy across the time series and participants. Error bar represents bootstrapped 95% C.I. **C**. Variations of SVM weights across the time series that correspond to the classifiers that were trained to decode exogenous versus endogenous cue validity. Topography indicates weights at the time point where we observed SVM peak values for ipsilateral and contralateral electrodes relative to the location of target onset.

### Electrophysiology: Decoding similar effects of exogenous and endogenous attention

We trained classifiers on one form of visuospatial attention and then tested their ability to generalize to the other form of visuospatial attention (Figure 2B). In this way, we wanted to verify if classifiers trained on exogenous attention generalized to endogenous attention, and vise-versa.

We first trained and validated linear SVM classifiers to decode exogenous cue validity (i.e., exogenous cue valid versus exogenous cue invalid), and then tested their performance to decode endogenous cue validity (i.e., endogenous cue valid versus endogenous cue invalid). This analysis revealed that the decoding of exogenous cue validity was above chance-level at target onset (i.e., −16ms) and lasted for the entire epoch (Figure 4A). During pre-processing of the EEG data, we relied on regression modelling based on the non-cueing condition to control for the sensory effects of the exogenous and endogenous cues and better isolate the effects of the target event. This result demonstrates that the classifiers were still able to capture ongoing sensory effects of the exogenous cue occurring shortly before the target event, despite correction (see Methods for details). The averaged peak decoding accuracy across participants for exogenous attention occurred at 223ms (95% CI, [123ms 323ms]) after target onset (Figure 4B). We examined the weights of the SVM and observed two peaks of SVM weights – a first one occurring before 200ms that corresponds to variations of electrodes on the contralateral side, and then a second one occurring after 200ms based on centroparietal and temporal fluctuations (Figure 4C). Importantly, we tested the generalization of these classifiers trained to decode the cue validity effects of exogenous attention on their ability to decode the cue validity effects of endogenous attention. We observed two significant clusters, a first one from 200 to 472ms, and a second from 564 to 612ms (Figure 4A). Furthermore, the averaged peak decoding accuracy for endogenous cue validity was 289ms (95% CI, [249ms 330ms]) after target onset (Figure 4B). These results corroborate the hypothesis that there are shared brain processes between exogenous and endogenous attention.

**Figure 4.**
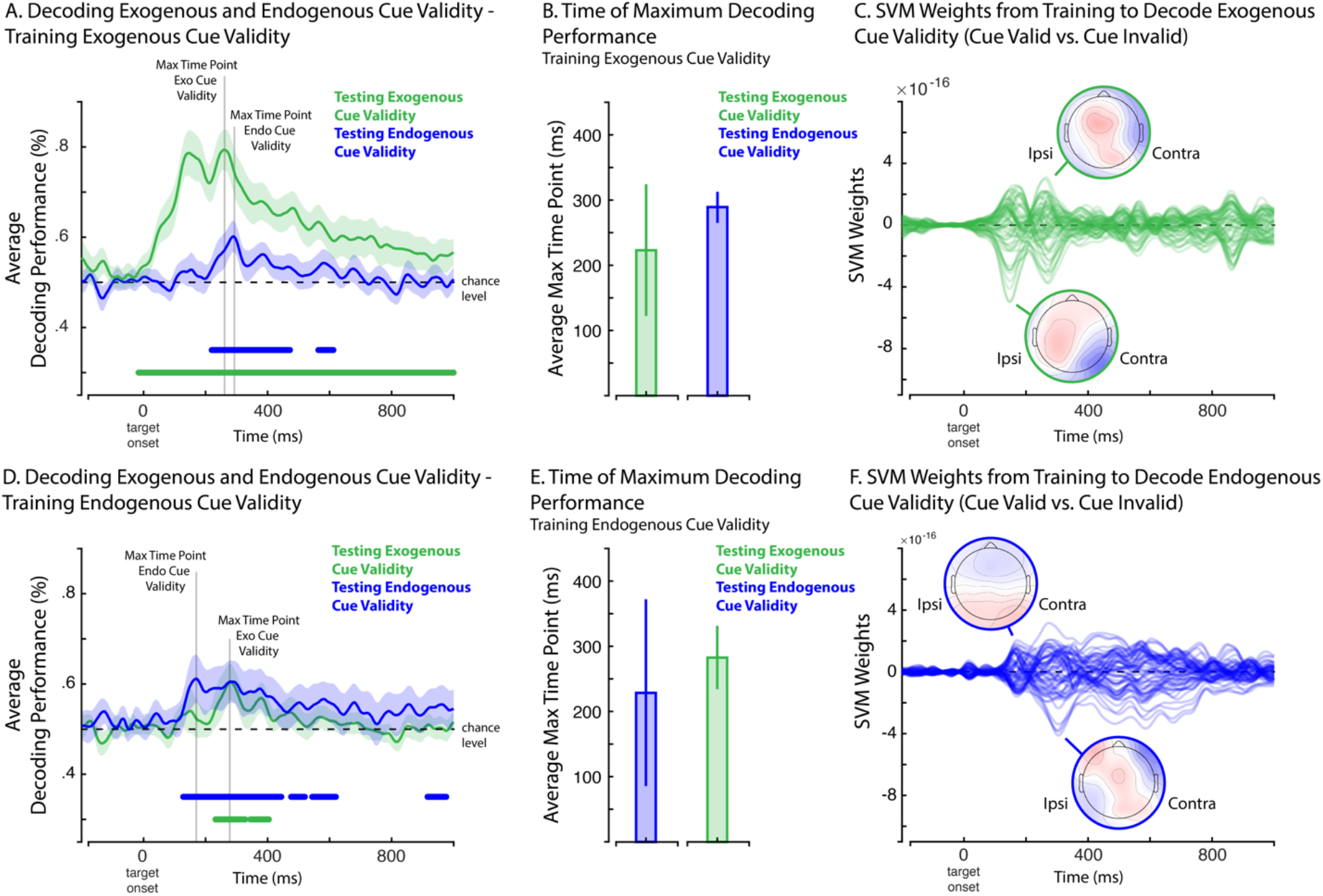
Group averages of accuracy from target-locked EEG traces for classifiers that were trained to decode exogenous cue validity (**A**; exogenous cue valid versus exogenous cue invalid) or endogenous cue validity (**D**; endogenous cue valid versus endogenous cue invalid) then evaluated on exogenous cue validity (green) and endogenous cue validity (blue). We used one sample t-tests and random cluster permutation to test for significant differences in classification performances against chance-level (50%) for both attention conditions. Horizontal lines indicate the temporal segments of significant differences in decoding accuracy for exogenous attention (green line) and endogenous attention (blue line). Shaded areas represent the 95% C.I. Averaged time point for maximal decoding accuracy across the time series and participants for classifiers that were trained for decoding exogenous cue validity (**B**) and endogenous cue validity (**E**), and then evaluated on exogenous cue validity (green bar graph) and endogenous cue validity (blue bar graph). Error bars represent bootstrapped 95% C.I. Variations of SVM weights across the time series that correspond to the classifiers that were trained to decode exogenous cue validity (**C**; exogenous cue valid versus exogenous cue invalid) and endogenous cue validity (**F**; endogenous cue valid versus endogenous cue invalid). Topography indicates weights at the time point where we observed SVM peak values for ipsilateral and contralateral sides relative to the location of target onset.

We used the same approach to evaluate whether SVM classifiers trained to decode endogenous cue validity (endogenous cue valid versus endogenous cue invalid) generalize to exogenous cue validity (exogenous cue valid versus exogenous cue invalid; Figure 2B). When we evaluated the performance of these classifiers for decoding endogenous cue validity against chance-level, we observed four significant clusters. A first one ranging from 128 to 444ms, the second one from 476 to 520ms, the third one ranging from 544 to 620ms, and lastly from 916 to 976ms (Figure 4D). Per our expectations, the effects of endogenous cue validity therefore emerged early after target onset. Here, the averaged time point for the maximal decoding accuracy across participants was 229ms (95% CI, [66ms 392ms]) after target onset (Figure 4E). The corresponding SVM weights comprised ipsilateral fluctuations at the posterior site before the 200ms mark relative to target onset followed by a more centro-frontal of weights fluctuations occurring beyond the 200ms mark post target-stimulus (Figure 4F). Importantly, we found two significant clusters when we applied this model to decode exogenous attention cue validity (exogenous cue valid versus exogenous cue invalid). A first one emerged from 232 and 328ms, and then a second one from 344 to 404ms (Figure 4D). The averaged time point of maximal decoding accuracy across participants for exogenous attention was 282ms (95% CI, [234ms 331ms]) post target stimulus (Figure 4E). Altogether, these results for models that are trained on endogenous attention and then tested on exogenous attention are consistent with the previous ones where we reported the outcome of SVM models that were trained on exogenous attention and then tested on endogenous attention. In both instances, the trained model generalized to other attention processing beyond 200ms post-stimulus.

We observed that transfer learning from exogenous attention to endogenous attention resulted in a slightly larger significant cluster than transfer learning from endogenous to exogenous attention. However, this difference is marginal: Comparing the decoding performance from these assessments did not reveal a significant difference between them. Indeed, one would expect that the information shared between exogenous and endogenous attention will be recovered across assessments. The difference in cluster size between these assessments is likely due to greater consistency of the exogenous attention neural response across trials, thereby improving the signal-to-noise ratio and hyperplane parameter estimations when training the linear SVM model in the context of exogenous attention. Greater consistency follows from the involuntary and effortless nature of the exogenous orienting response (MacLean et al., 2009).

We further compared the performance of SVM classifiers trained to decode exogenous attention and tested on exogenous attention against that of classifiers trained to decode endogenous and tested on endogenous attention. Here, the performance for decoding exogenous attention was greater than for decoding endogenous attention from 36 to 584ms after target onset (Appendix A).

### Early ERP components relate to perceptual facilitation

Our previous results uncovered early (<200ms post-target onset) differential and later overlapping effects between exogenous and endogenous cue validity. Here, we aimed to explore whether these effects contribute to perceptual facilitation (i.e., faster response times for cued trials relative to uncued ones). We relied on single-trial ERPs and mediation analyses to test this hypothesis. Our approach was similar to previous techniques in single-trial ERPs analysis (e.g., Nunez et al., 2017). To obtain single-trial waveforms, we first applied principal component analysis (PCA) on the target-locked averaged waveforms of the non-cueing condition across the entire scalp (see Methods). This allowed us to uncover components pertaining to visual processing in the absence of explicit orientation of attention. From this analysis we retained three components that explained 90.92% of the total variance (i.e., 41.98% for the first component, 36.69% for the second component and 12.25% for the third and last one; see Appendix B). The loading coefficients of the first component revealed a pattern that encompassed the posterior region that was skewed towards the ipsilateral side, the second component also showed a posterior topographical pattern, albeit on the contralateral side, and the third one exhibited a more centroparietal topographical pattern (Figure 5A; Appendix B). Next, we used the PCA weights obtained from the non-cueing condition to project the single-trial target-locked EEG data from the exogenous and endogenous spatial cueing conditions (Appendix B). This resulted in 3 sets of single-trial loadings for each attention condition. Averaged waveforms for exogenous and endogenous attention (i.e., cue valid and cue invalid trials) across these different components are presented in the Appendix C. See the Method section for details.

**Figure 5.**
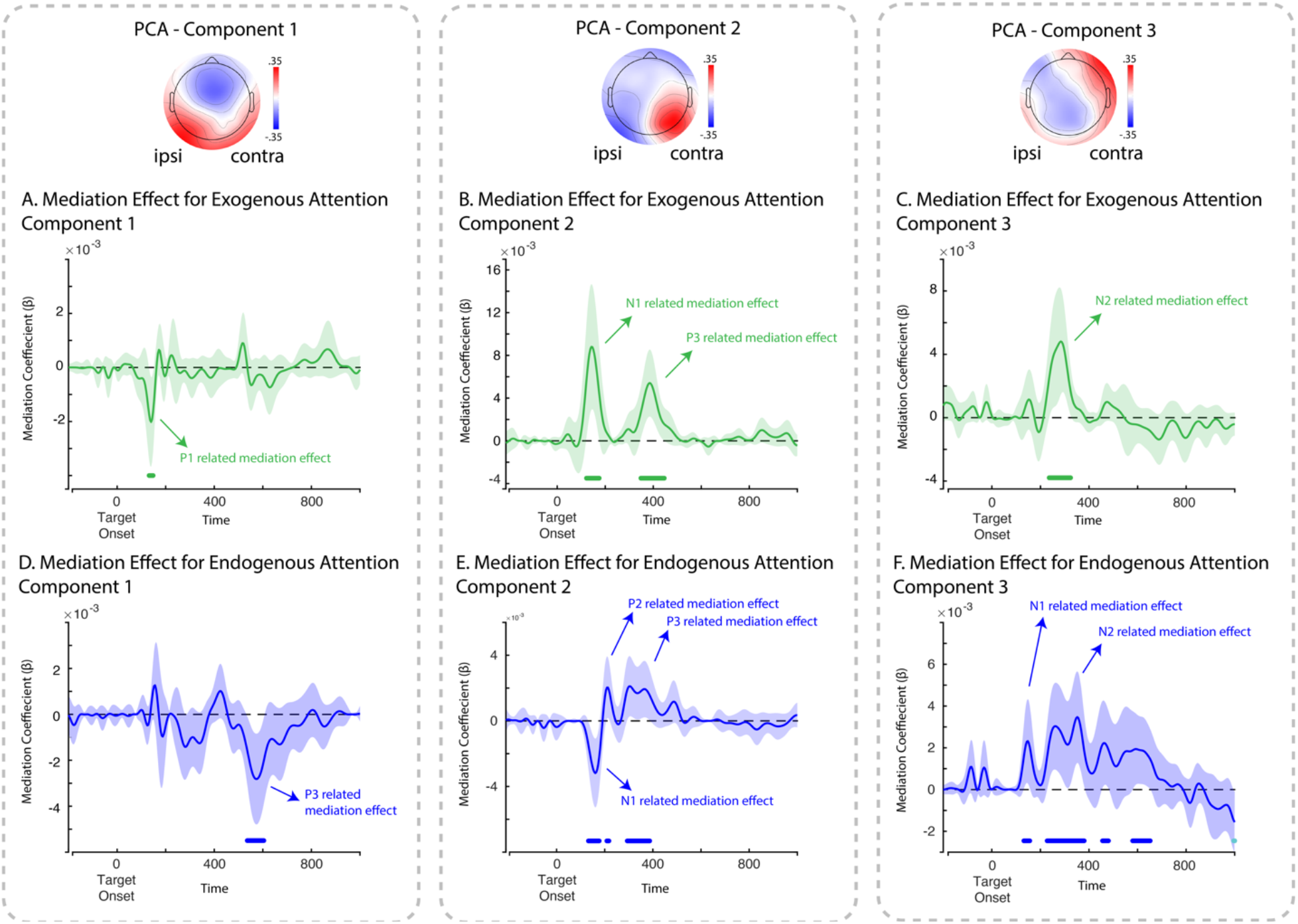
Single-trial ERPs mediation effects for exogenous (green) and endogenous (blue) across the three components we isolated applying PCA to the non-cueing condition. Coefficients show parameter estimates for mediation effects involving the ERPs across all three PCA components. Green and blue bars represent significant effects for cluster-level alpha < .05. Shaded areas represent 95% C.I. Topographies for the first, second, and third component for target-related contra- and ipsilateral are shown at the top.

We applied mediation analyses separately across the 3 components. Here, we evaluated whether single-trial ERPs mediate the relationship between cue validity and response times based on hierarchical regression analyses. Meditation coefficients were evaluated across the time series and corrected for multiple comparisons using cluster-based permutation tests. See Appendices D and E for coefficients of the a, b, c’ paths. For exogenous attention, a significant cluster emerged between 132 and 148ms post-target for the first component (R^2^_mediation_ = 12.58%; Figure 5A) that indicated a partial mediation effect (Appendix D shows path c’ coefficients). This effect was related to increased amplitude of the P1 for exogenous cued trials versus uncued ones (the coefficients for the a path are presented in Appendix D). For the second component, we observed two significant clusters – a first one between 132 and 184ms that corresponded to decreased amplitude of the N1 for cued trials versus uncued ones (R^2^_mediation_ = 12.51%), and a second one from 356 to 452ms which reflected decreased amplitude of the P3 (R^2^_mediation_ = 12.52%; Figure 5B). Both clusters indicated a partial mediation (see Appendix for D for c’ path coefficients). Lastly, we observed a significant cluster indicating a partial mediation for the third component from 236 to 324ms (R^2^_mediation_ = 12.53%; Figure 5C), which matched decreased amplitude of the N2.

In our last analysis, we applied the same mediation procedure for endogenous attention. We found a significant cluster from 536 to 604ms that indicated a partial mediation, which corresponded to increased amplitude for the P3 (R^2^_mediation_ = 13.53%; Figure 5D; we present a and c’ paths coefficients in Appendix E). Moreover, for the second component, we observed two early significant clusters from 140 to 184ms that matched an increased N1 (R^2^_mediation_ = 13.54%) and a much smaller one from 216 to 224ms that corresponded to an increased P2 (R^2^_mediation_ = 13.5%; Figure 5E; a path coefficients in Appendix E); and a later cluster that corresponded to an increased P3 amplitude between 300 and 392ms after target onset (R^2^_mediation_ = 13.5%; Figure 4D). Lastly, we observed four significant clusters for the third component. A first cluster that corresponded to decreased amplitude for the N1 from 132 to 156ms post-target onset (R^2^_mediation_ = 13.5%), a second one that corresponded to decreased amplitude of the N2 from 228 to 380ms post-target onset (R^2^_mediation_ = 13.49%), and then two clusters (i.e., from 456 to 480ms and from 456 to 652ms post-target onset) that reflected a lasting difference in amplitude between endogenous cued and uncued trials.

## Discussion

In the present work, we investigated how exogenous and endogenous attention achieve perceptual facilitation through distinct and similar neural processes. We relied on an easy target discrimination task where the magnitude of cueing benefits for exogenous and endogenous attention was comparable across both systems, as evidenced by Bayes factor analyses (Figure 1). We then leveraged MVPA over target-locked ERPs to investigate when and where exogenous and endogenous attention neural processes are different and similar (Figure 2). Overall, our results indicate that exogenous and endogenous attention differ early on (<200ms post-target onset) in terms of their neural processes. In contrast, both attention systems share similar neural processes later (∼300ms after target onset). Lastly, we linked the observed differential and overlapping effects between attention systems to their perceptual facilitation using a single-trial ERPs meditation analysis. Altogether, our findings demonstrate how endogenous and exogenous facilitation effects are governed by overlapping and differential processes, further supporting the multifaceted view of visuospatial attention.

Our findings show that exogenous and endogenous attention differed along the entire time series. This difference was maximal around 170ms (Figure 3A and 3B). In accordance with our hypothesis, differential effects between exogenous and endogenous attention therefore occurred early after target onset. These findings dovetail previous work that highlights early (between 100 and 200ms after target onset) differences between exogenous and endogenous attention (Hopfinger & West, 2006). We similarly observed that the maximal difference between exogenous and endogenous attention was most prominent on the contralateral topographical posterior site relative to the stimulus’ location (Figure 3C). These attention processes, therefore, operate on separate stages of visual processing on the contralateral side.

We also observed important differences in decoding accuracies between exogenous and endogenous cue validity (Figure 4A, Figure 4D, and Appendix A). This outcome denotes a larger effect size for exogenous cue validity compared to endogenous cue validity– a difference we can readily observe for the components of our PCA (Appendix C). Here, the difference in amplitude between valid and invalid trials was greater for exogenous attention across the P1, N1 and N2, which correspond to the period when maximal classification was achieved for discriminating exogenous cue validity from endogenous cue validity. Therefore, while the cueing effects of exogenous and endogenous attention were similar at the behavioral level, exogenous attention yielded greater modulation of the sensory signal at the neural level. This result likely reflects the prominence of bottom-up processes that drive the perceptual benefits of exogenous attention (Fernández & Carrasco, 2020), which produced a superior cueing effect in the EEG. Conversely, recent evidence links the locus of top-down processes in endogenous attention to the frontal eye field (Fernández, Hanning, & Carrasco, 2023), which seems to have produced smaller cueing effects in the EEG.

We should also note that the cueing of exogenous attention followed from a peripheral sensory stimulation occurring closely in time to the target event. These experimental factors likely improved the decoding accuracy of exogenous attention despite our regression approach to remove cue-related contamination from target-related processing – thus, our regression approach did not fully control for this contamination. The observation that classification of exogenous cue validity was already above chance-level shortly before target onset aligns with this interpretation. Nevertheless, as we detailed in the previous paragraph, larger cueing effects for exogenous attention better accounts for our decoding results.

In line with the MPVA results, our mediation modelling approach indicates that these effects occurred over the P1-N1 complex, which is consistent with previous work in the field (Hopfinger & Mangun, 1998, 2001; Hopfinger & West, 2006). For this contralateral component, exogenous attention increased the amplitude of the P1 and decreased that of the N1, as evidenced by the values of the coefficient for the a path along the time series, whereas endogenous attention did not alter the amplitude of the P1 and increased the amplitude of the N1 (Appendices C, D and E). These outcomes exhibit different patterns between them. Prevailing views relate modulations of these early ERPs to sensory gain for attended events relative to unattended ones (Hillyard et al., 1998; Itthipuripat et al., 2022; M. M. Müller et al., 2006). Accordingly, our results imply that exogenous and endogenous attention boost the sensory signal through different means: Exogenous attention appears to influence sensory processing earlier than endogenous attention. This outcome is consistent with findings from functional magnetic resonance imaging that show differential effects for exogenous and endogenous attention at the level of the visual areas (Dugué et al., 2020; N. G. Müller & Ebeling, 2008).

Importantly, our mediation modelling approach shows that early differential effects between attention processes relate to perceptual facilitation (Figure 5). Here, we found that exogenous and endogenous attention partly mediated perceptual facilitation by modulating differently the components we isolated using PCA. For our first component, which exhibited an ipsilateral topographical pattern at the visual level, we observed increased amplitude of the P1 following exogenous attention partly mediated the relationship between the exogenous attention cue and perceptual facilitation (Figure 5A). In contrast, we did not observe any such effect for early ERP following endogenous attention. Instead, endogenous attention contributed to facilitation via modulations of the P3 (Figure 5D), which aligns with previous work showing a dominance of endogenous attention at later stages (Hopfinger & West, 2006). In this way, we observed a distinct pattern for the first component: An early mediation effect for exogenous attention and late one for endogenous attention, which means that differences between exogenous and endogenous attention also occur later during processing.

Differential effects also emerged for the contralateral EEG component of our PCA (i.e., our second component) where both exogenous and endogenous produced partial mediation effects through opposite patterns. Whereas the mediation of exogenous attention involved decreased amplitude of the N1, meditation of endogenous attention occurred through increased amplitude of the N1. This pattern further underscores how differential effects contribute to perceptual facilitation. Note that modulation of exogenous attention on the amplitude of the contralateral P1 did not contribute to this mediation effect. Accordingly, the influence of exogenous attention over the P1, a neurophysiological marker of exogenous attention (Martín-Arévalo et al., 2016), partially mediated perceptual facilitation for the ipsilateral topographical component only, not the contralateral one. While the maximal difference between exogenous and endogenous attention occurred at the contralateral site early after target onset, our results show that exogenous attention facilitates perception through a more ipsilateral component, not the contralateral one. Lastly, we also observed a partial mediation effect related to decreased amplitude of the N1 following endogenous attention for the third and last EEG component (Figure 3F). Therefore, endogenous attention partly facilitates perception by boosting the N1 at the contralateral site and decreasing the amplitude of the N1 at centroparietal level. Altogether, these findings emphasize that facilitation effects for exogenous and endogenous attention emerge through different stages of visual processing (i.e., boosting the ipsilateral P1 for exogenous attention and the ipsilateral P3 as well as the contralateral N1 for endogenous attention). Modulations of the P1, an early ERP component, for exogenous attention likely reflect the stimulus-driven nature of this attention process, whereby the salient features of the peripheral cue trigger early bottom-up visual processes that enhance the processing of upcoming event at the same location (Chica et al., 2013). In turn, modulations of the N1, a later component, following endogenous attention likely reflect the top-down nature of this attention process, which encompass feedback re-entrant loops from the frontoparietal circuits that enhance neural activity in the sensory areas in a goal-driven manner (Carrasco, 2011).

Our findings also emphasize similarities between exogenous and endogenous during visual processing. We observed shared neural processes beyond the period of the P1-N1 complex (Figure 4A, 4B, 4D and 4E). For both exogenous and endogenous attention. The time point of maximal similarities between attention systems corresponded to a more central topography (Figure 4C and 4F). These results denote that beyond early differential effects, exogenous and endogenous impact sensory processing in a similar fashion. These results support growing literature suggesting that these attention processes share neural processes at the sensory level (Landry et al., 2022). Likewise, our mediation analysis also shows analogous effects for the second and third component (which corresponded to a contralateral component and a more centroparietal topographical pattern, respectively; see Appendix B). Specifically, looking at the values of the coefficients for the a path, both exogenous and endogenous attention yielded smaller amplitudes of the P3 over the contralateral component and of the N2 for the centroparietal one (see Appendices C, D and E). These results add to a body of mixed evidence that reports similar (Martín-Arévalo et al., 2016) and differential effects of exogenous and endogenous attention on the P3 (Hopfinger & West, 2006). Such heterogeneity in the literature brings about the possibility that this may reflect task-specific effects. Still, few studies directly compare exogenous and endogenous in the same experimental context. Our findings therefore advance the field by showing analogous effects between both exogenous and endogenous attention for later stages of visual processing (>200ms post-target) in the same experimental context. Our results, similarly, nuances the idea that later stages of visual processing are dominated by one form of attention, and instead highlight how both systems impact these later stages in an analogous fashion.

In addition to early differential effects, perceptual facilitation also involves overlapping modulations for later stages of visual processing (>200ms post-target onset; Figure 5B, 5C, 5E and 5F). This analogous effect, however, is driven by the processing during uncued target events. Indeed, we observed greater amplitude for later ERPs during invalid trials for both exogenous and endogenous attention, while these effects partly mediated perceptual facilitation. It is possible that similarities between exogenous and endogenous attention follows from the re-orientation of attention resources towards uncued target events (Corbetta, Patel, & Shulman, 2008), which would ultimately modulate the later stages of sensory processing in a similar fashion. This interpretation entails that changes in the amplitude of the N2 and the P3 relates to the processing of unexpected events (e.g., Bocquillon et al., 2014; Debener, Makeig, Delorme, & Engel, 2005). Both the exogenous and the endogenous cues, therefore, may have generated expectations about the target’s location, even if the exogenous cue was non-predictive of the target’s location and participants were made aware of this fact. In sum, the advent of the target event at the uncued location could therefore represent an unexpected event that recruits later stages of target-related processing following the re-orientation of attention resources. On the other hand, these effects could also reflect the recruitment of greater resources for target-related processing to recover from the cost of having to re-orient perceptual resources. In our study, both the N2 and the P3 were predictors of response times regardless of attention and cueing effects. Modulations of these ERPS, therefore, reflect performance during target discrimination. In short, the re-orientation of attention resources encompasses greater recruitment of these target-related resources.

Although we uncovered how distinct and shared processes between exogenous and endogenous attention facilitate perception, our findings revealed a partial mediation, which entails that additional neural processes contribute to the cueing effect. In this regard, one may expect that changes in alpha oscillations would likewise contribute to this effect (Peylo, Hilla, & Sauseng, 2021). Moreover, it also remains unclear how distinct and shared processes between exogenous and endogenous attention relate to specific computations in shaping their effects (e.g., Jigo, Heeger, & Carrasco, 2021).

We also note that, since differential effects occurred early after target onset and lasted the entire time series (Figure 3A) and overlapping effects encompassed only part of the time series and emerged later (Figure 4A and 4D), these effects occurred in parallel. This outcome is hardly unexpected. The effects of exogenous and endogenous attention on visual processing encompass multiple components, as demonstrated by the present results and the large body of work we previously highlighted. Given that some of these components parallel each other in time, differential and overlapping effects between exogenous and endogenous attention can therefore co-exist. In the same vein, our mediation analyses show that both exogenous and endogenous influence the same component, albeit to a different extent, which also yields differential effects. Hence, the effects of exogenous and endogenous attention on visual processing are multifaceted and involves separatable and shared features.

Combining the strengths of MVPA, single-trial ERPs and mediation modelling, our study delivers a comprehensive assessment of the distinct and overlapping neural processes of exogenous and endogenous attention. Our findings align with the dichotomous account of visuospatial attention by showing that early differential effects contribute to shaping perceptual facilitation (Carrasco, 2011; Chica et al., 2013). They also align with reports that highlight differences between exogenous and endogenous attention at the neural level such as the frontoparietal network (e.g., Bowling, Friston, & Hopfinger, 2020), the temporoparietal junction (e.g., Dugué, Merriam, Heeger, & Carrasco, 2018), and the visual areas (e.g., Dugué et al., 2020). Critically, MVPA enabled us to highlight how these attention processes also yield similar effects further along the visual stream. This result is consistent with the notion of shared neural processes between them. In this regard, we found that both differential and overlapping effects contributed to shaping perceptual facilitation. Specifically, our findings imply that exogenous and endogenous attention exert different effects on early sensory processing, which include the P1-N1 waveforms, and then impact processes more closely related to perceptual decisions making in a similar fashion, as indexed by the N2 and P3 waveforms. In sum, our findings are consistent with the idea that exogenous and endogenous attention hardly reduce to a single neural marker and involve multiple stages of sensory processing (Martín-Arévalo et al., 2016).

## Acknowledgments

M.L. & J.D.C. acknowledges fellowships from the Natural Science and Engineering Research Council of Canada. K.J. is supported by funding from the Canada Research Chairs program (950-232368) and a Discovery Grant from the Natural Sciences and Engineering Research Council of Canada (2021-03426), a Strategic Research Clusters Program (2023-RS6-309472) from the Fonds de recherche du Québec – Nature et technologies, and an IVADO-Apogée fundamental research project grant. Lastly, we would like to thank members of the Razlab who helped with data collection.

## Contributions

M.L. and J.D.C. designed, implemented the study, and analyzed the data. K.J. provided feedback on data analysis and interpretation of results. All authors wrote the manuscript.

## Data availability

Data and code supporting the current study is publicly available on the Open Science Framework repository: https://osf.io/43fyv/?view_only=5990b42982ff4c958f35056a49bcf624

## Code Availability

Code for the analyses supporting the current study is publicly available on the Open Science Framework repository:

## Appendix A

**Figure A1.**
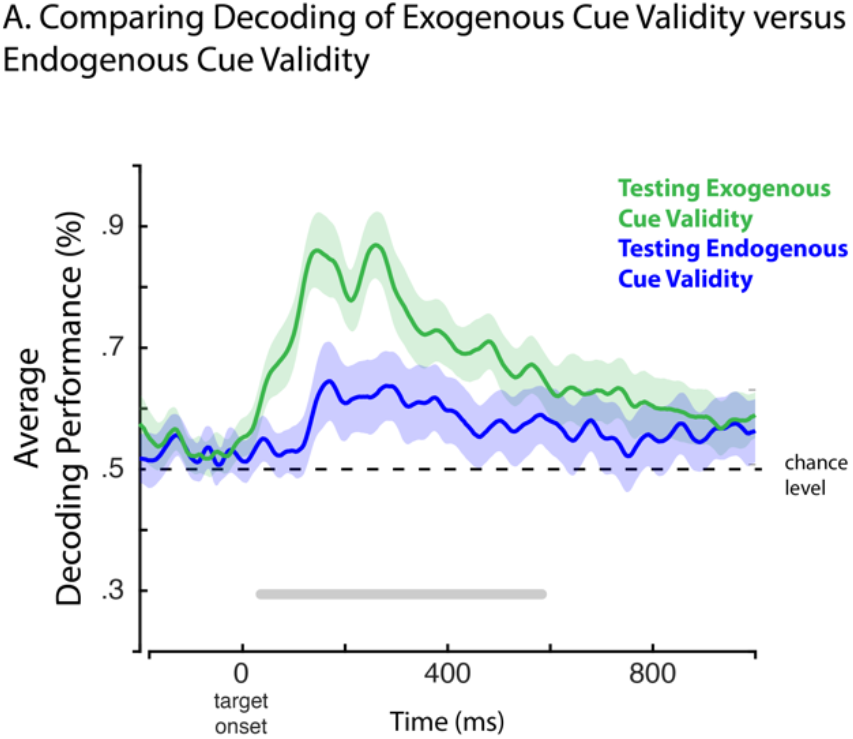
**A**. Group averages of decoding accuracy from target-locked EEG traces for exogenous cue validity and endogenous cue validity. We used pairwise t-tests and random cluster permutation to test for significant differences in classification performances. Horizontal grey line indicates the temporal segments of significant differences in decoding accuracy. Shaded areas represent the 95% C.I.

## Appendix B

**Figure B1.**
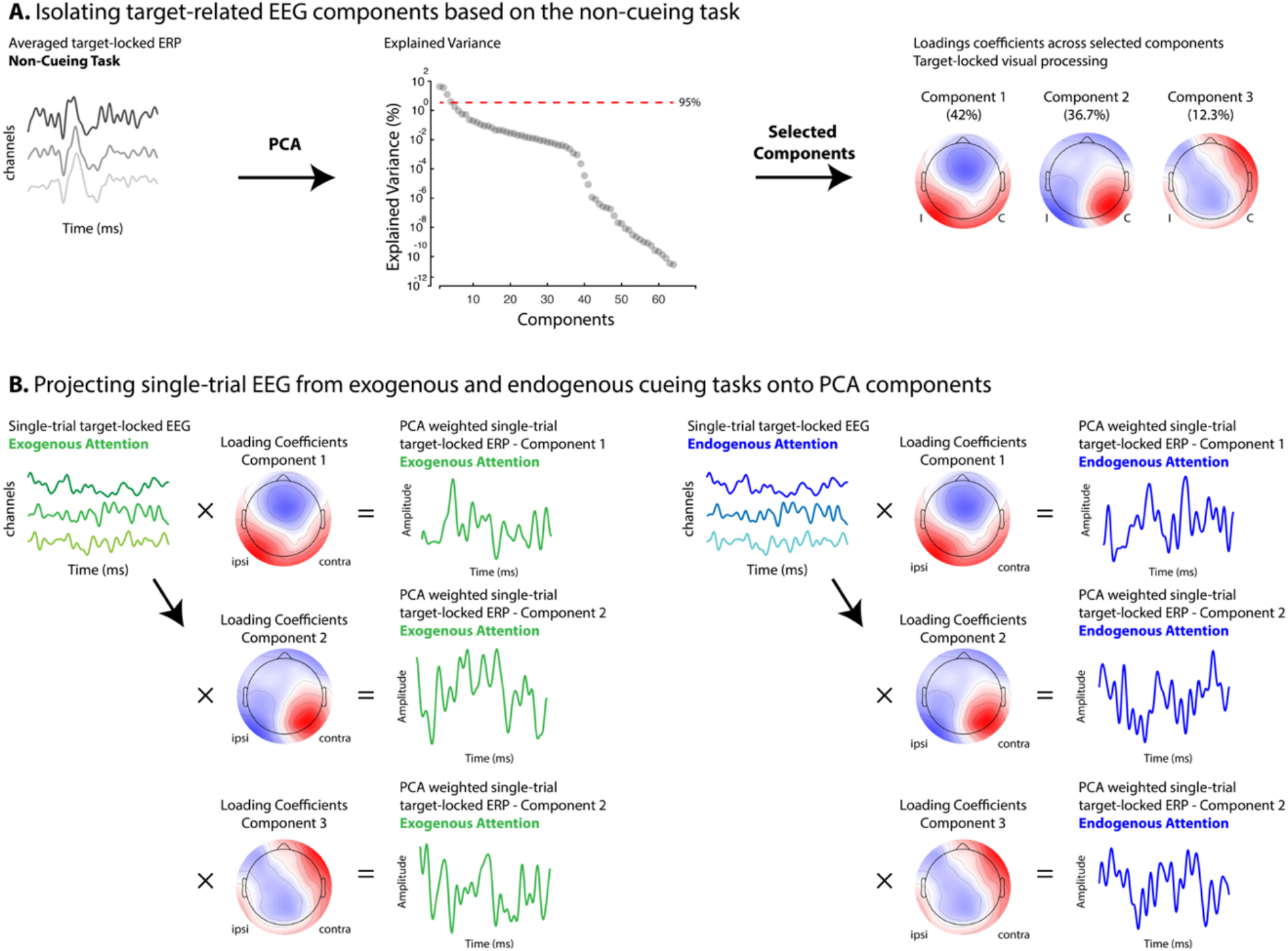
We applied principal component analysis (PCA) to target-locked waveforms (time by channels) from the non-cueing task averaged across participants to identify the components related to visual processing (**A**) and extracted 3 components that explained 90.92% of the variance. We performed random permutations to identify our components of interest based on a 95% threshold. Next, we projected single-trial EEG (time by channels) from the exogenous and endogenous cueing tasks onto each component (vector of loading coefficients) separately (**B**). We obtained PCA weighted single-trial ERPs for each component across exogenous and endogenous attention.

## Appendix C

**Figure C1.**
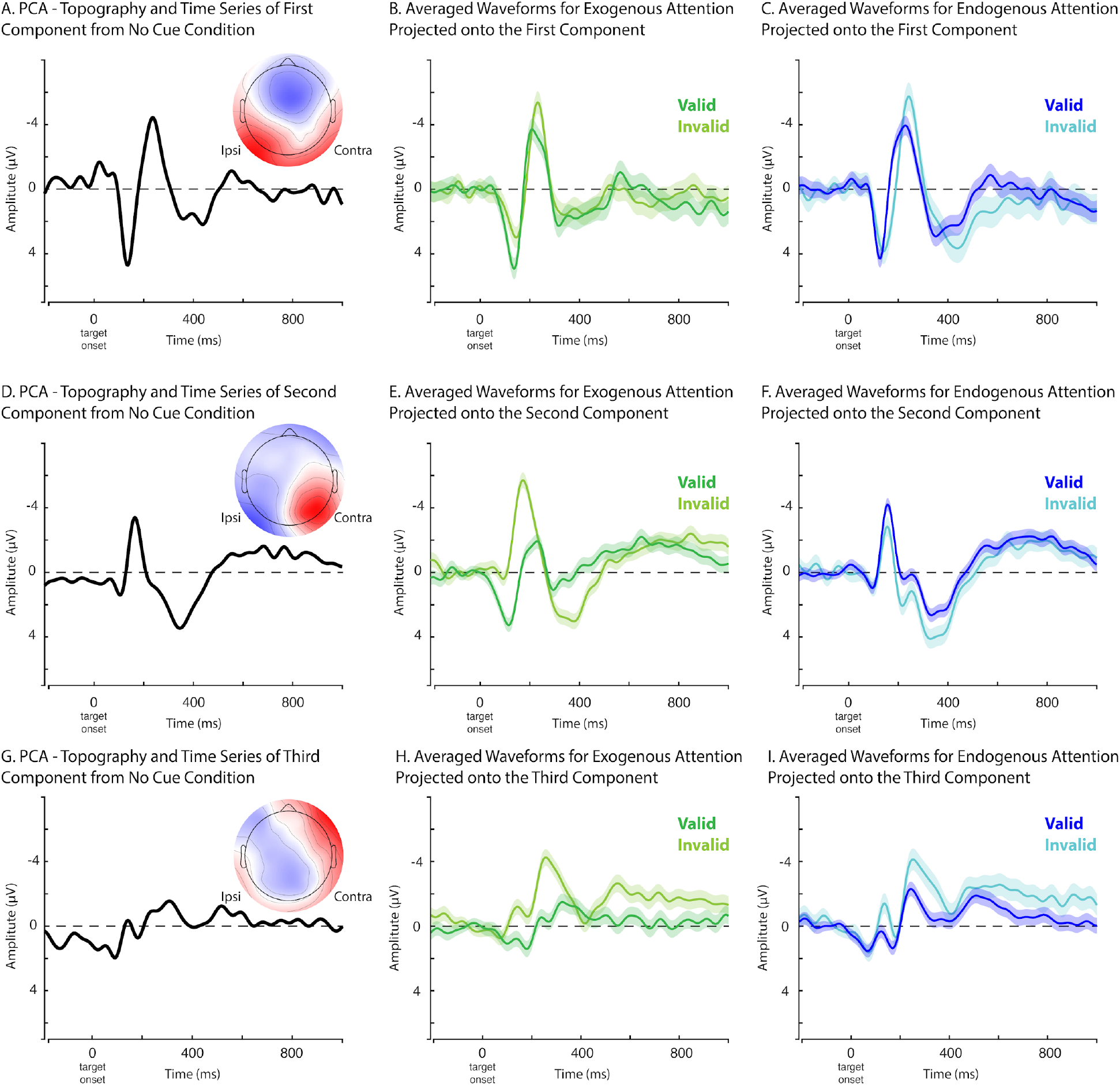
Time series and topography of first (A), second (D) and third (G) components from PCA applied to the target-locked averaged waveform from the non-cueing condition. Target-locked grand averaged ERPs from all trials for exogenous and endogenous cue valid and cue invalid projected onto the first (B,C), second (E,F), and third (H,I) component. Waveforms for exogenous and endogenous cue validity include 95% C.I.

## Appendix D

**Figure D1.**
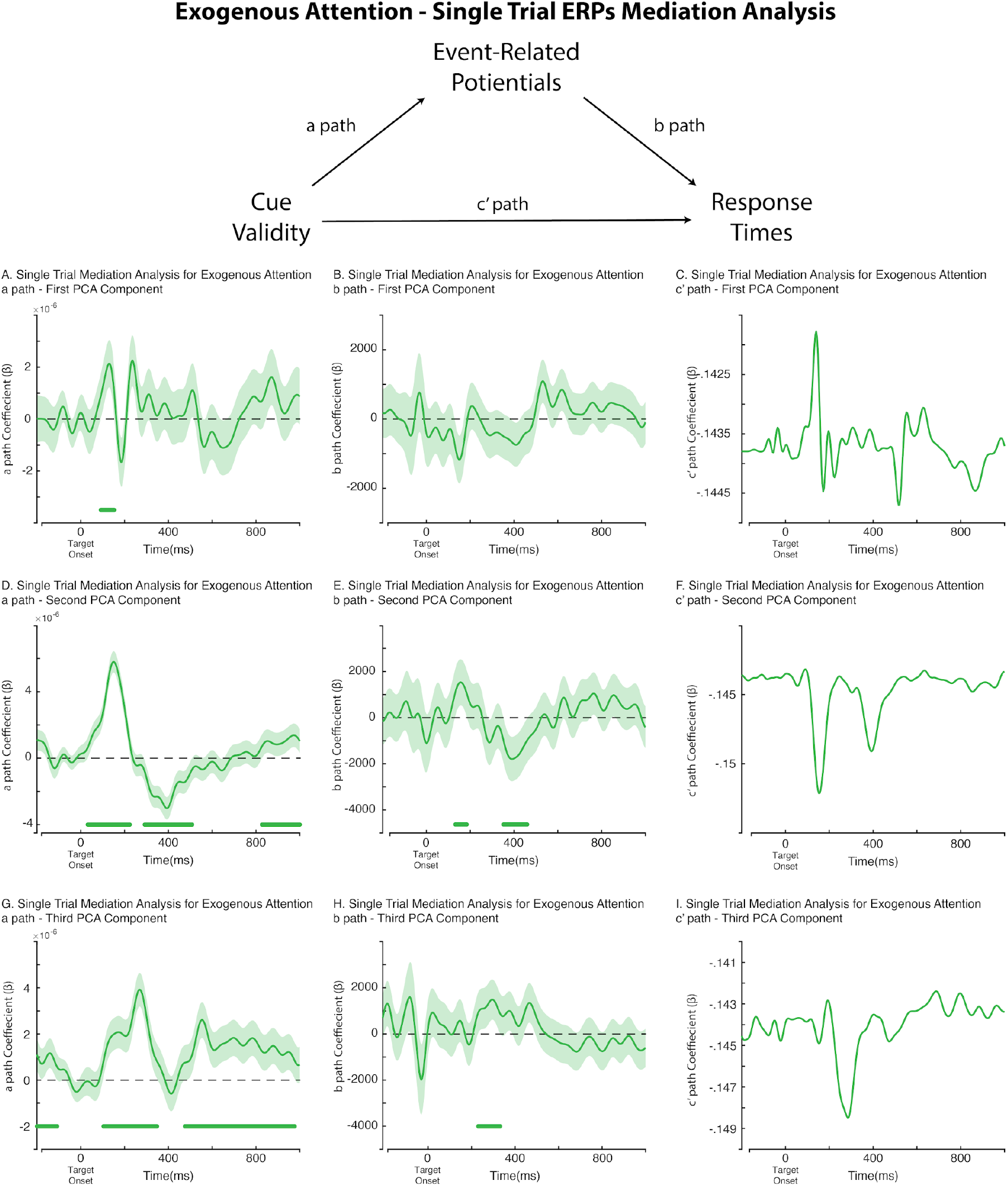
Time series of a path, b path and c’ path for single trial ERPs mediation analysis where ERPs partly mediate the relationship between exogenous cue validity and response times. Mediation analysis was done across first (A,B,C), second (D,E,F) and third (G,H,I) components from PCA applied to the target-locked averaged waveform from the non-cueing condition. Waveforms for a and b paths encompass 95% C.I. for mediation analysis coefficients.

## Appendix E

**Figure E1.**
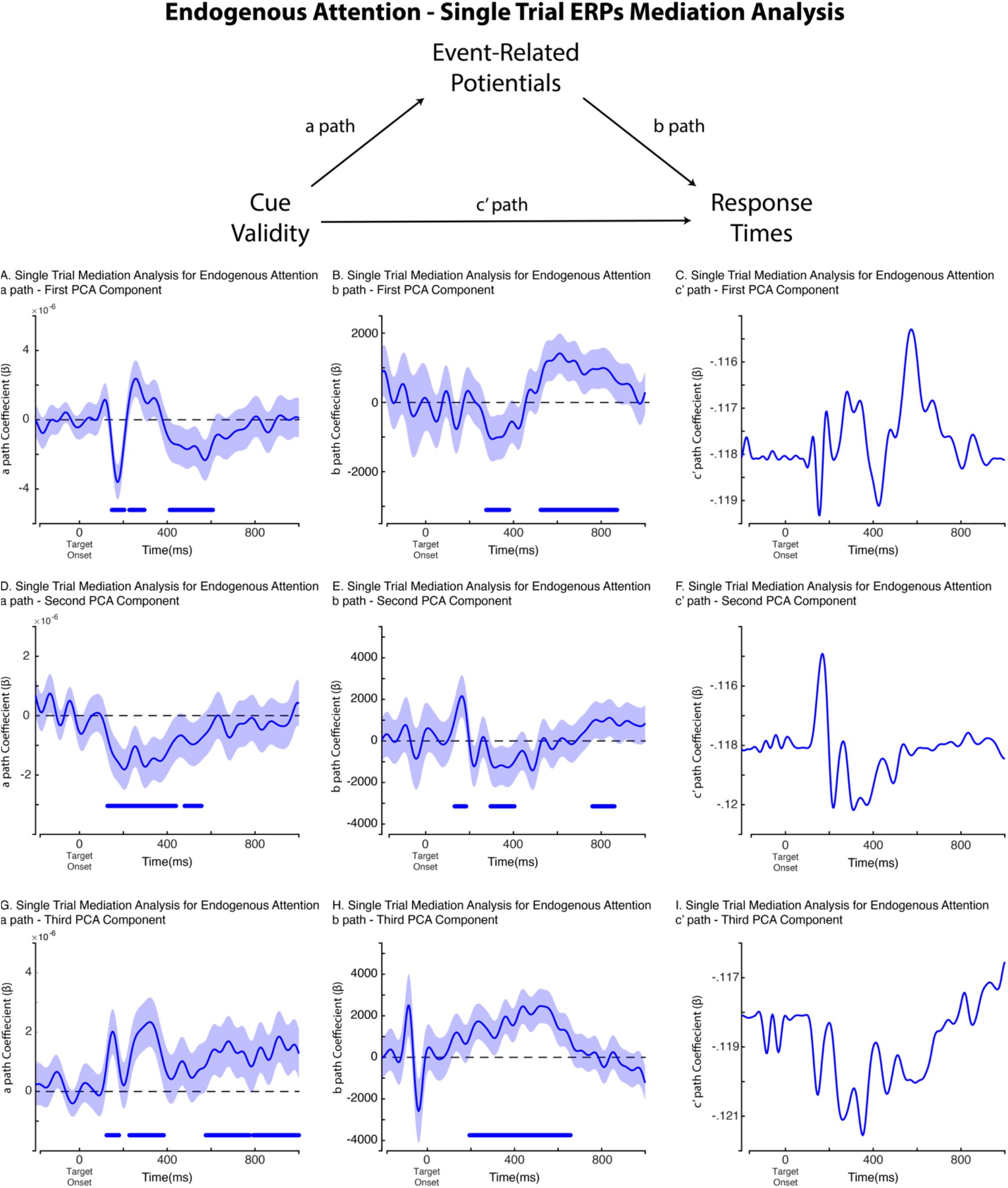
Time series of a path, b path and c’ path for single trial ERPs mediation analysis where ERPs partly mediate the relationship between endogenous cue validity and response times. Mediation analysis was done across first (A,B,C), second (D,E,F) and third (G,H,I) components from PCA applied to the target-locked averaged waveform from the non-cueing condition. Waveforms for a and b paths encompass 95% C.I. for mediation analysis coefficients.

